# A phylogenomically informed five-order system for the closest relatives of land plants

**DOI:** 10.1101/2022.07.06.499032

**Authors:** Sebastian Hess, Shelby K. Williams, Anna Busch, Iker Irisarri, Charles F. Delwiche, Sophie de Vries, Tatyana Darienko, Andrew J. Roger, John M. Archibald, Henrik Buschmann, Klaus von Schwartzenberg, Jan de Vries

## Abstract

The evolution of streptophytes had a profound impact on life on Earth. They brought forth those photosynthetic eukaryotes that today dominate the macroscopic flora: the land plants (Embryophyta) [1]. There is convincing evidence that the unicellular/filamentous Zygnematophyceae—and not the morphologically more elaborate Coleochaetophyceae or Charophyceae—are the closest algal relatives of land plants [2, 3, 4, 5, 6]. Despite the species richness (>4,000), wide distribution, and key evolutionary position of the zygnematophytes, their internal phylogeny remains largely unresolved [7, 8]. There are also putative zygnematophytes with interesting body plan modifications (e.g., filamentous growth) whose phylogenetic affiliations remain unknown. Here, we studied a filamentous green alga (strain MZCH580) from an Austrian peat bog with central or parietal chloroplasts that lack discernible pyrenoids. It represents *Mougeotiopsis calospora* PALLA, an enigmatic alga that was described more than 120 years ago [9], but never subjected to molecular analyses. We generated transcriptomic data of *M. calospora* strain MZCH580, and conducted comprehensive phylogenomic analyses (326 nuclear loci) for 46 taxonomically diverse zygnematophytes. Strain MZCH580 falls in a deep-branching zygnematophycean clade together with some unicellular species, and thus represents a formerly unknown zygnematophycean lineage with filamentous growth. Our well-supported phylogenomic tree lets us propose a new five-order system for the Zygnematophyceae, and provides evidence for at least five independent origins of true filamentous growth in the closest algal relatives of land plants. This phylogeny provides a robust and comprehensive framework for performing comparative analyses and inferring the evolution of cellular traits and body plans in the closest relatives of land plants.

## RESULTS AND DISCUSSION

### Morphology and phylogenetic position of a filamentous zygnematophyte without pyrenoids

Strain MZCH580 forms unbranched filaments with smooth cell walls and rounded tips (Figure 1A, B). Infolded cross walls (‘replicate walls’) or rhizoids known from some filamentous zygnematophytes [10] were not observed in our cultures. The filaments of strain MZCH580 tend to fragment as the cultures age, but cells divide and grow back into new filaments when fresh medium is added (Figure 1C, D). Interphase cells are 10-15 μm wide (mean = 12 μm, n = 40) and 12-55 μm long (mean = 22 μm, n = 80), and usually contain a single chloroplast. The chloroplast lacks visible pyrenoids and has a variable shape ranging from an off-centre straight plate (Figure 1D) to a more parietal morphology, like a channel or half-pipe (Figure 1A, B). The 3D reconstruction of confocal fluorescence data reveals a common intermediate morphology (Figure 1E). The lateral sides of half-pipe-shaped chloroplasts display clear indentations, which are rare in filamentous green algae with chloroplasts of similar morphology (Figure 1A, B, arrows)— *Entransia fimbriata* (Klebsormidiophyceae), for example, has fimbriate or lobed chloroplasts, but of much more irregular morphology [11]. The nucleus is spherical (4-6 μm in diameter, n = 40) with a prominent central nucleolus (1-3 μm in diameter, n = 40), and always closely associated with the chloroplast (Figure 1A, D; nuc). Both chloroplast and nucleus are surrounded by a thin sheath of cytoplasm and opposed to or surrounded by a large vacuole (Figure 1D; asterisks).

**Figure 1:**
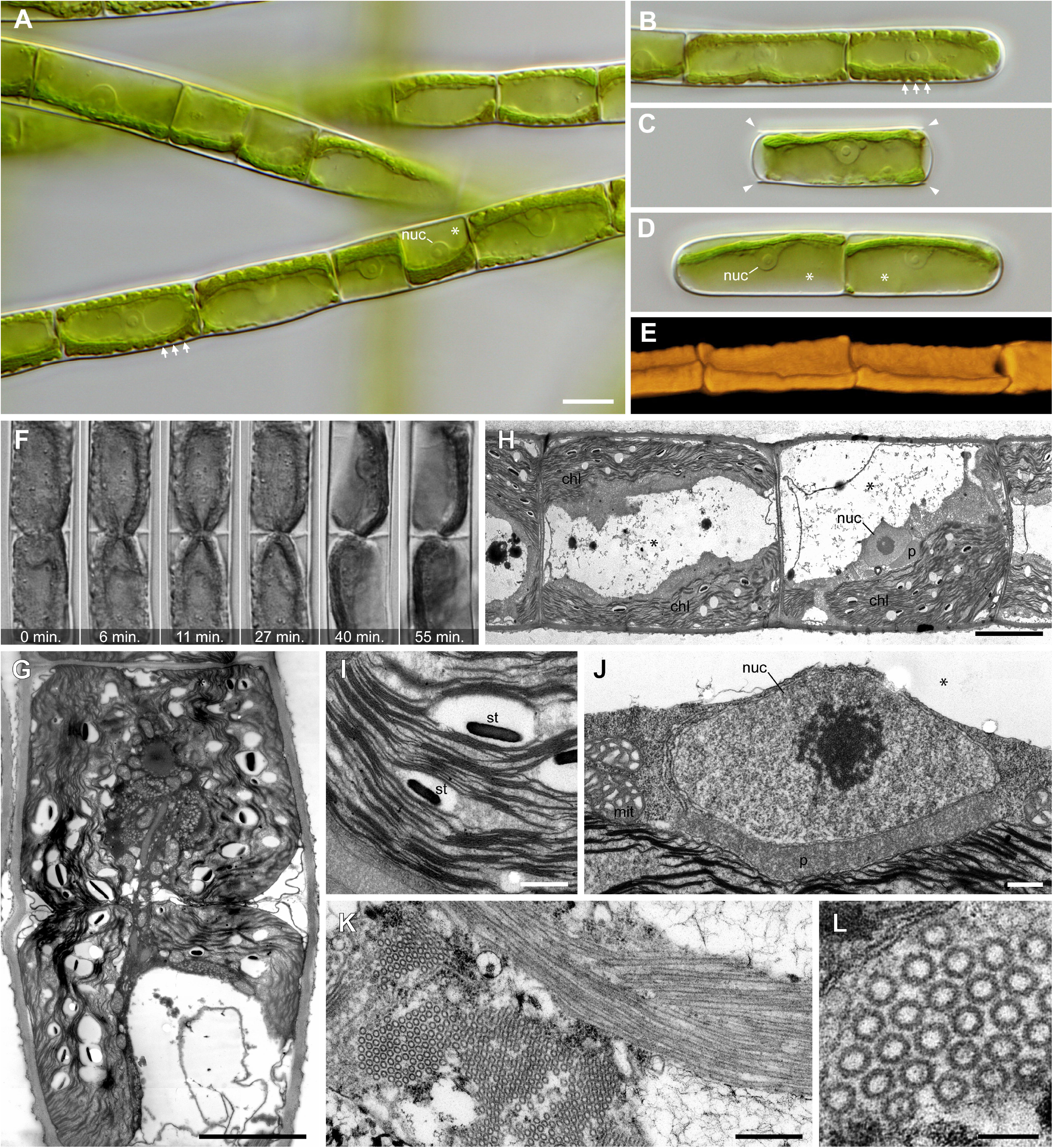
Morphology, cell division and ultrastructure of Mougeotiopsis calospora strain MZCH580. **A:** Filaments with cells of varying length; DIC. Note the indented chloroplast margins (arrows), the prominent nuclei (nuc) and the large vacuoles (asterisk). **B:** Filament with rounded tip; DIC. **C:** Singe cell after fragmentation with cell wall remnants (arrowheads); DIC. **D:** Two-celled filament with smooth tips; DIC. Note the prominent nuclei (nuc) and the large vacuoles (asterisks). **E:** Three-dimensional reconstruction of the chloroplasts based on their autofluorescence; confocal microscopy. **F:** Time series of a dividing cell shows ingrowing cross wall; DIC. **G:** Ultrathin section through a dividing cell reveals the ingrowing cell wall (see plasma membrane) and the chloroplast in division. **H:** Ultrathin section through vegetative filament showing the position of the nucleus (nuc), peroxisome (p), chloroplasts (chl) and vacuoles (asterisks). **I:** Ultrathin section of starch grains between the thylakoids of the chloroplast. **J:** Ultrathin section of the nucleus (nuc) with nucleolus, the large, elongate peroxisome (p), and mitochondria (mit). The vacuolar space is marked by the asterisk. **K:** Ultrathin section of bundled macrotubules in cross section (left) and longitudinal section (right). **L:** Detail of macrotubules in cross section. Scale bars 10 μm in A (applies also for B–D); 5 μm in G and H; 500 nm in I–K; 100 nm in L. See also Figure S1.

Cell division is intercalary and involves the centripetal formation of a cross wall (Figure 1F; Videos S1 and S2). We did not observe any phragmoplast-like structure as known from many streptophyte algae [12, 13, 14]. Instead, ingrowing cell wall material seemed to pinch off the chloroplast (Figure 1F, Videos S1 and S2), which is corroborated on the ultrastructural level (Figure 1G). It appears that the chloroplast does not divide before the inset of cytokinesis and that the cell division in strain MZCH580 largely depends on furrowing (cleavage, thus centripetal cell wall ingrowth). However, we cannot exclude the existence of a phragmoplast and our ultrastructural data of late stages of cytokinesis seem compatible with phragmoplast-like structures as known from many streptophyte algae including other zygnematophytes (e.g. *Spirogyra, Mougeotia* [12, 13]). Furthermore, our ultrastructural data confirm that the chloroplasts of strain MZCH580 lack pyrenoids, but contain numerous lentiform starch grains (up to ∼1 μm) interspersed between the thylakoids (Figure 1H, I). Other noteworthy ultrastructural characteristics of strain MZCH580 include a giant peroxisome situated between the nucleus and the chloroplast (Figure 1J), and the occurrence of macrotubules (∼44 nm in diameter) in cells under unfavorable conditions (Figure 1K, L). A single peroxisome of similar localization was also reported for the filamentous algae *Klebsormidium* (Klebsormidiophyceae) [15, 16, 17] and *Zygogonium* (Zygnematophyceae) [18], suggesting that this is a rather widespread character in streptophyte algae. However, the filamentous zygnematophytes *Mougeotia, Spirogyra* and *Zygnema* contain numerous, much smaller peroxisomes, which do not exceed 1 μm in our TEM sections (Figure S1).

Based on taxonomic comparisons (see Taxonomic Appendix for details), we apply the name *Mougeotiopsis calospora* to strain MZCH580. However, as we did not observe any sexual processes (conjugation, flagellated gametes), zoospores or aplanospores in our cultivated material, the suspected affinity to the zygnematophytes remained uncertain. While analysis of the *rbcL* gene (coding for the large chain of ribulose-1,5-bisphosphate carboxylase/oxygenase) placed strain MZCH580 within the streptophytes, a robust phylogenetic placement was not possible. To scrutinize the phylogenetic position of strain MZCH580, we generated RNAseq data by Illumina sequencing and performed a *de novo* transcriptome assembly with Trinity. The resulting transcriptome has a completeness of 96.3% assessed with BUSCO (benchmarked universal 272 single-copy orthologs) and contains 52,188 predicted open reading frames (ORFs). We built a comprehensive multigene dataset of 326 conserved proteins (see Methods) from streptophyte algae, land plants, and select chlorophyte algae as outgroup, with 84 taxa in total (see species and deposited data in Methods). Our phylogenomic inferences with a sophisticated site-heterogeneous model of protein sequence evolution (LG+PMSF(C60)+F+Γ) resulted in a well-supported phylogeny, whose overall topology is in line with current knowledge about streptophyte evolution (cf. Figure S1 and [6]). Strain MZCH580 groups within the Zygnematophyceae with full nonparametric bootstrap support, and forms a deep-branching lineage with the unicellular *Serritaenia* sp. (strain CCAC 0155) and “*Mesotaenium endlicherianum*” (strain SAG 12.97). Hence, strain MZCH580, referred to as *Mougeotiopsis calospora hereafter*, is clearly distinct from other filamentous genera (*Mougeotia, Spirogyra, Zygnema and Zygnemopsis*), and represents a new lineage of zygnematophytes with filamentous growth.

### Phylogenomics support a five-order taxonomy of the Zygnematophyceae

Previous phylogenies based on single (or few) marker genes have suggested that the traditional taxonomic separation into the two orders Desmidiales and Zygnematales does not reflect the evolutionary relationships of the Zygnematophyceae [7, 8]. And yet the taxonomy of this important algal class remains unresolved, in part due to the lack of robust phylogenetic data. Our multigene phylogeny clearly demonstrates that the Zygnematales as previously defined (all filamentous members plus unicells that are not placoderm desmids) are paraphyletic. Instead, the Zygnematophyceae comprise at least five deep-branching clades that we feel can be treated at the level of orders (Figure 2).

**Figure 2:**
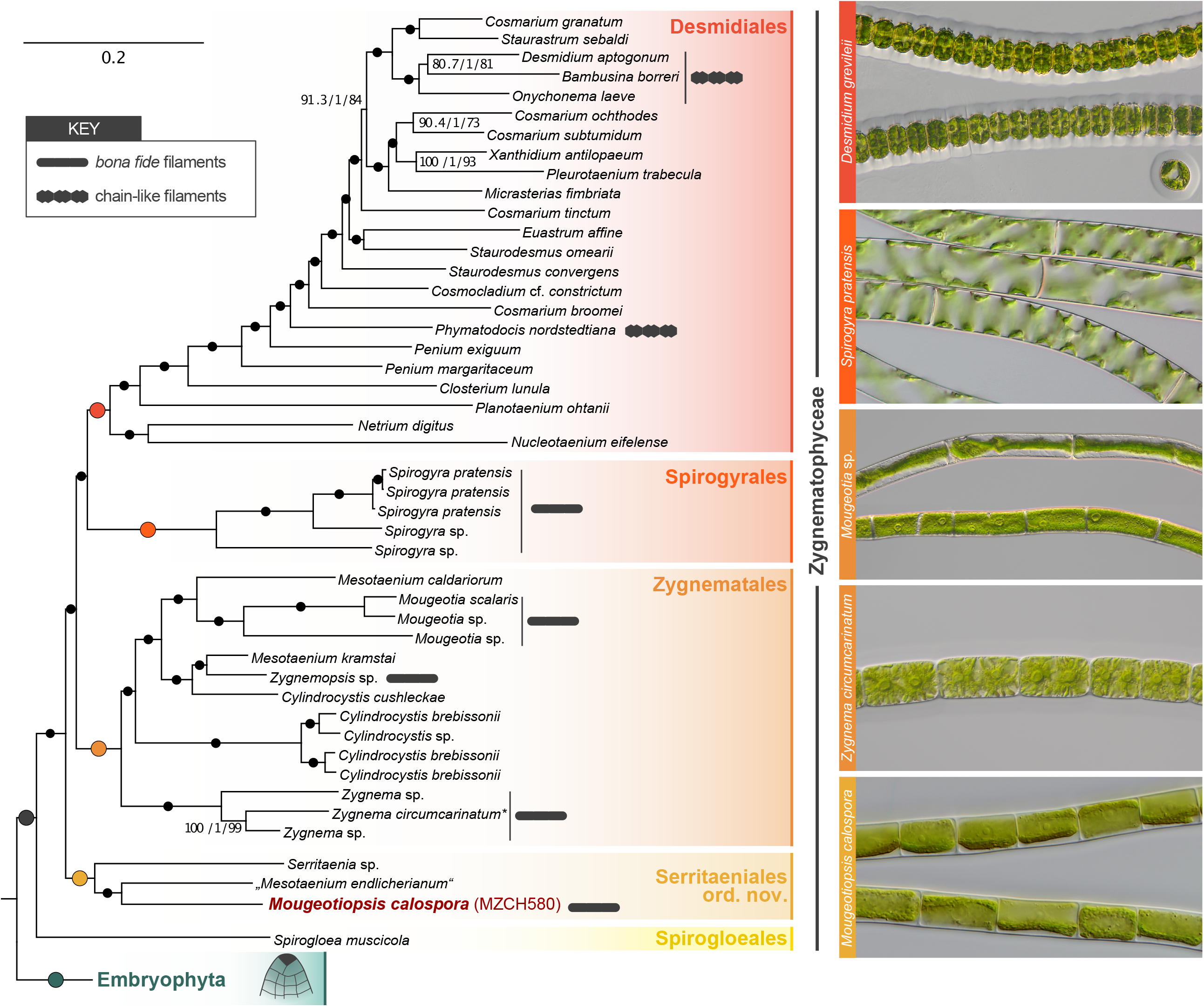
Position of strain MZCH580 in a well-resolved zygnematophycean phylogeny based on 326 genes. Section of the phylogenomic tree limited to zygnematophytes and embryophytes. Support values from three analyses (SH-aLRT/aBayes/nonparametric bootstrapping) are shown at the corresponding branches, except for branches with maximum support (marked by black dots). The Zygnematophyceae comprise five deep-branching clades, which are here defined as orders. Gray symbols highlight zygnematophytes that form chain-like filaments (see *micrograph* of *Desmidium*) and *bona fide* filaments (see micrographs of *Spirogyra, Mougeotia, Zygnema* and *Mougeotiopsis*). Scale bar for phylogeny is 0.2 substitutions per site. The entire phylogenomic tree with all streptophyte taxa is shown in Figure S2. Asterisk: a recent study by Feng et al. [80] found that SAG698-1a might be Z. *cylindricum* instead of Z. *circumcarinatum*.

We introduce a new, phylogenomically informed five-order taxonomy of the Zygnematophyceae, by reinterpreting existing ordinal names and introducing a new order for *Mougeotiopsis* and its unicellular relatives (see Taxonomic appendix). The Serritaeniales *ord. nov*. currently comprises the name-giving genus Serritaenia (unicells with a plate-like chloroplast and a mostly aerophytic life style; [19]), the genome-sequenced strain SAG 12.97 (often referred to as “*Mesotaenium endlicherianum*” [20]; unicells with half-pipe-like chloroplasts and an aquatic life style) and *Mougeotiopsis calospora*, strain MZCH580. Although these species differ markedly in growth form (unicells vs. filaments), their chloroplasts are all characterized by indented or undulated margins [19,20] that are otherwise rare in zygnematophytes. Yet, *Mougeotiopsis calospora* is the only known zygnematophyte that lacks pyrenoids.

Our data corroborate the position of the Spirogloeales, consisting of the unicellular *Spirogloea muscicola* (formerly *Spirotaenia muscicola*), as sister lineage to all other Zygnematophyceae [20]. For the remaining part of the phylogenomic tree, we redefine three traditional orders. The Zygnematales are now limited to a morphologically diverse clade comprising unicellular zygnematophytes currently assigned to *Cylindrocystis* and *Mesotaenium* plus three distinct branches of filamentous members (*Mougeotia, Zygnema* and *Zygnemopsis*); the recovered topology demonstrates the polyphyly of the unicellular genera belonging to that order (*Cylindrocystis, Mesotaenium*), which require a taxonomic revision in the future. Chloroplasts of the Zygnematales are either stellate (*Cylindrocystis, Zygnema, Zygnemopsis*) or ribbon/plate-like with smooth margins (*Mesotaenium, Mougeotia*).

The *Spirogyra* species with their characteristic helical chloroplasts form another, deep-branching clade, which is here defined as Spirogyrales Clements, 1909 (Figure 2 and Taxonomic appendix). This order was initially introduced to comprise algae of yellow-green appearance (including *Spirogyra*) and some fungal families [21]. We limit the concept of the Spirogyrales to those zygnematophycean algae that form the sister clade of the Desmidiales in our phylogeny. The latter order mainly comprises symmetric unicells with a pronounced central constriction (isthmus) and ornamented cell walls. However, at the base of the clade containing these typical placoderm desmids are three genera (*Netrium, Nucleotaenium, Planotaenium*), which display a much simpler morphology (no cell wall ornamentations, no isthmus) and were formerly classified with the Zygnematales (in the family Mesotaeniaceae) [22]. Interestingly, the same arrangement was previously recovered by combined analyses of three genes (nuclear SSU rRNA, *rbcL*, chloroplast LSU rRNA) [23], and is here confirmed by phylogenomics. It appears that the desmids with elaborate cell shapes and complex cell walls (e.g. *Cosmarium, Penium, Micrasterias, Xanthidium*) descended from unicellular ancestors with a simpler structure. Hence, the genera *Netrium, Nucleotaenium* and *Planotaenium* are here formally included in the order Desmidiales. The internal phylogeny and taxonomy of the Desmidiales, however, needs to be resolved by extended taxon sampling in the future, as many classically recognized desmid genera (e.g. *Cosmarium, Penium, Staurodesmus*) are not monophyletic.

### On the unicellularity of the ancestral zygnematophyte

Our robust phylogenetic framework of the zygnematophytes now enables comparisons of species in an evolutionary context, and assessment of evolutionary scenarios with greater confidence. It is remarkable that the majority of zygnematophycean species are unicellular [24], as most of their streptophyte relatives (Embryophyta, Coleochaetophyceae, Charophyceae, Klebsormidiophyceae, Chlorokybophyceae) display some kind of multicellularity, from sarcinoids to three-dimensional tissues [25]. However, some zygnematophycean lineages exhibit more developmental complexity such as the formation of filaments, sometimes even with rhizoids or branched cells [25, 26]. Traditionally these filamentous members have been bundled in the family Zygnemataceae [22], but a close relationship of them was not recovered in previous phylogenies [7, 8].

Our fully-supported phylogenomic tree reveals at least five separate lineages that contain true filaments, found in three orders (Figure 2): *Spirogyra* (Spirogyrales), *Mougeotia, Zygnema, Zygnemopsis* (all Zygnematales), and *Mougeotiopsis* (Serritaeniales). Other filamentous taxa (e.g. *Temnogametum iztacalense, Zygogonium ericetorum*) await genomic/transcriptomic sequencing and phylogenomic placement [27, 28]. The cells of all these filamentous species have straight and relatively simple cell walls, no central constrictions, and display an intimate cell-cell contact (i.e. typical cross walls)—yet without plasmodesmata [29]. At the same time, there are also filamentous desmids (e.g. *Desmidium, Bambusina, Onychonema*, and *Phymatodocis*), which differ markedly from the aforementioned lineages in their cellular details and filament morphology. The cells of *Desmidium, Bambusina, Onychonema* and *Phymatodocis* display the typical characters of desmid cells (e.g. central constriction, cell wall ornamentation) and rather appear as cell chains. Together with the fact that the filamentous desmids are nested within the unicellular desmids, it is conceivable that there are distinct types of filamentous growth in the Zygnematophyceae, which evolved independently; we account for this possibility in our analyses (see Figure 3 and below). Given the paucity of unicellular life forms in the streptophytes, previous studies noted that unicellular zygnematophytes might have arisen by reductive evolution from a morphologically more complex ancestor [5, 30, 31, 32, 33]. Based on our current phylogeny, this seems less parsimonious than a unicellular ancestor and multiple independent origins of filamentous growth in the zygnematophytes. It would require at least seven losses of multicellularity versus five independent origins of filamentous growth (if one excludes the filamentous desmids).

**Figure 3:**
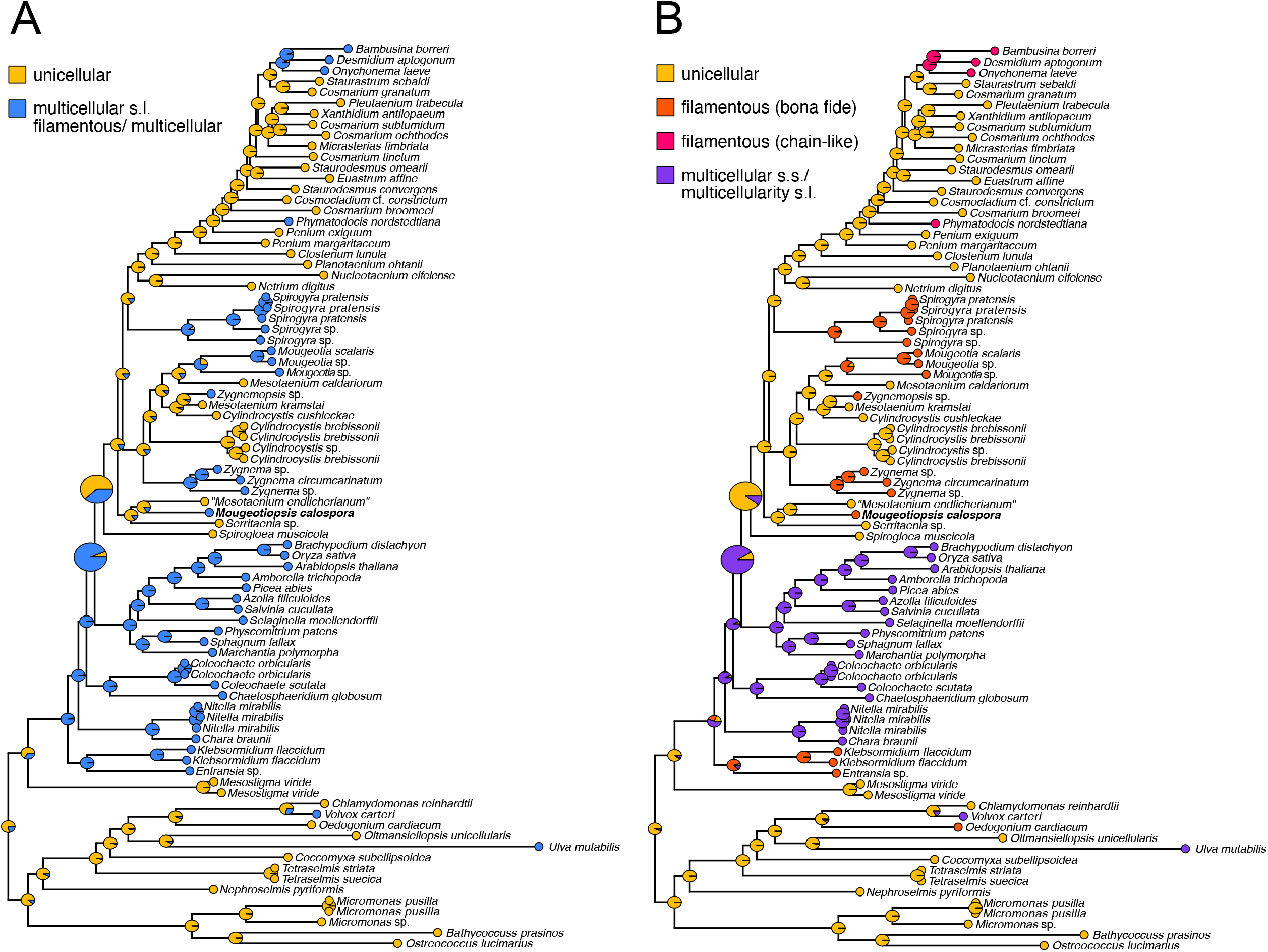
Ancestral character state reconstruction for unicellular or multicellular (including filamentous) growth characters. Growth types were coded as either a simplistic (A) two- and more nuanced (B) four-character state distributions to reflect different levels of complexity regarding the possibilities/hypotheses for the homology of growth types: yellow = unicellular, blue = multicellular *sensu lato* (including filamentous growth), orange = *bona fide* filamentous growth, pink = chain-like filaments (desmids), and purple = multicellular growth *sensu stricto*.

In an attempt to infer the body plan of the common ancestor of zygnematophytes, we performed ancestral character state reconstructions (ACSR) with various data coding strategies concerning the types of multicellularity (Figure 3). Irrespective of how the growth types were coded, a unicellular zygnematophyte ancestor was consistently inferred by our analyses, albeit with varying support (PP=0.58-0.93). Hence, we infer up to five tentative independent origins of true filamentous growth, and two additional independent origins of chain-like filaments (in the Desmidiales) (unicellular ancestors have PP=0.80-1.00); under this scenario, the last common ancestor of the Zygnematophyceae and land plants was likely filamentous or multicellular (PP=0.91-0.93), whereas the last common ancestor of Zygnematophyceae was likely unicellular (PP=0.58-0.89). Given the effect of character coding in these analyses, we conclude that expanding our knowledge about the homology of the various types of multicellular and filamentous body plans in the green algae is essential.

Filamentous growth as observed in the Zygnematophyceae can be considered the least elaborate type of multicellularity [34]. Yet, the cellular and molecular traits underpinning this growth type remain obscure. The multiple growth type transitions in the zygnematophytes are consistent with parallel evolution from a common molecular machinery, but the relative simplicity of filamentous growth renders convergent evolution equally plausible. The hypothetical unicellular lynchpin at the base of the Zygnematophyceae is an attractive hypothesis: it could explain why zygnematophytes lack plasmodesmata (see e.g., [29]), why the cross walls often look distinct from other streptophytes, and perhaps even why the group as such returned to a cleavage-like cell division mechanism (see [14]). Future research on the different filamentous lineages will need to establish a deeper understanding of the molecular machinery underpinning their common morphology.

In addition, recent culture-based efforts to explore terrestrial zygnematophytes indicated a high diversity of unicellular lineages [35], which are not yet covered by genomic/transcriptomic sequencing and might change the evolutionary picture. Biased taxon sampling is indeed a serious problem for ACSR [36, 37] and thus genomic sequencing of further zygnematophytes is an important task for the future. Given the great age of the lineages involved and the sparse fossil record, important information might be obscured by extinction events as well; new discoveries of living or fossil taxa could easily lead to new interpretations. For now, our phylogenomic data demonstrate that the zygnematophytes comprise multiple transitions of their body plan, and enable the selection of relevant species for comparative cell biological research.

## Conclusion

The identification of the Zygnematophyceae as the sister lineage to land plants was surprising, in part because of their relatively simple body plans. The study of zygnematophycean trait evolution is a challenge because of their species richness, diverse morphologies, and unresolved phylogeny. We have provided a phylogenomic backbone and a congruent classification system for the closest algal relatives of land plants. Looking at algal growth types through the lens of phylogenomics reveals dynamic emergence and formation of filamentous and unicellular growth among the Zygnematophyceae—traits whose evolutionary history might also feature reductive evolution from a more complex ancestor of Zygnematophyceae and land plants.

## Taxonomic appendix

### Rationale for the application of the name Mougeotiopsis calospora to strain MZCH580

In terms of its gross morphology, strain MZCH580 resembles members of the genera *Klebsormidium* (Klebsormidiophyceae), *Ulothrix* (Ulvophyceae) and *Mougeotia* (Zygnematophyceae), all of which form unbranched filaments and have plate-like or parietal plastids. However, the absence of pyrenoids in strain MZCH580 is a major distinguishing character, as algae from the three mentioned genera (and classes) typically have prominent pyrenoids surrounded by a sheath of starch grains. There are, however, two historical descriptions from the late 19^th^ century that describe pyrenoid-lacking, filamentous green algae with plate like chloroplasts: *Mougeotiopsis calospora* Palla, 1894 and *Mesogerron fluitans* Brand, 1899. *Mougeotiopsis* is a putative zygnematophyte, as scalariform conjugation and the formation of zygospores was clearly documented [9]. Instead, *Mesogerron* was only described on the basis of vegetative material, and first suspected to be related to *Ulothrix* (Ulvophyceae, Chlorophyta). Based on the marked resemblance in their vegetative characters (filament width of 15-18 μm, cell architecture, and chloroplast morphology), *Mougeotiopsis* and *Mesogerron* were later treated as heterotypic synonyms (Krieger, 1941 [77]). Strain MZCH580 matches both descriptions concerning the varying cell length (including cells that are shorter than wide), cell architecture (plastid-associated nucleus) and chloroplast morphology (plate-like to parietal with pronounced lateral indentations), but it has somewhat smaller cells (filament width of 10-15 μm). The morphological similarity, however, is compelling, and variation in filament width is known for many closely related strains or species of green algae. We were unable to locate the type material of *Mougeotiopsis calospora*, but studied original material of *Mesogerron fluitans* (collected by F. Brand in 1899 and provided by the Herbarium of the Academy of Natural Sciences of Philadelphia). The dried filaments of that species were morphologically similar to those of strain MZCH580, especially in the marked variation in cell length observed in the filaments (Figure S3). Amplification of genetic material from this sample did not work.

### Rationale for establishing a new order, Serritaeniales ord. nov

In our phylogeny, the branch in question comprises three distinct groups of organisms: *Mougeotiopsis calospora* (one strain known), the genus *Serritaenia* (several strains known; [19]), and strain SAG 12.97, a unicellular zygnematophyte that is often referred to as “*Mesotaenium endlicherianum*”. Currently, there is only one existent ordinal name that is based on the mentioned taxon names, namely Mesotaeniales Fritsch. However, the phylogenetic position of the genus *Mesotaenium* is still uncertain, as the designation of strain SAG 12.97 is potentially based on misidentification. In the opinion of some authors (S.H. and A.B.), the morphology of SAG 12.97 does not conform with the description of the type species *M. endlicherianum* Nägeli. This problem was already recognised by other specialists for zygnematophycean algae who studied strain SAG 12.97 [78, 79]. Hence, we are hesitant to reuse the name Mesotaeniales and instead introduce a new ordinal name based on the well-studied and credible genus *Serritaenia*. Descriptions of the zygnematophycean orders defined in this study follow.

### Order Serritaeniales

S. Hess & J. de Vries ord. nov.

#### Diagnosis

comprises unicells and filaments with smooth sidewalls, cells with axial or parietal chloroplasts and simple cell walls (no pores and ornamentations), phylogenetically closely related to the type species (*Serritaenia testaceovaginata; rbcL* MW159377).

#### Type family (designated here)

Serritaeniaceae S. Hess & J. de Vries fam. nov.

#### Comment

Currently this order includes the genera *Serritaenia* and *Mougeotiopsis*.

### Family Serritaeniaceae

S. Hess & J. de Vries fam. nov.

#### Diagnosis

with characteristics of order Serritaeniales; unicells and filaments with smooth sidewalls, cells with axial or parietal chloroplasts and simple cell walls (no pores and ornamentations), embedded or not in the mucilage.

#### Type genus (designated here)

*Serritaenia* A.Busch & S.Hess 2021.

With the type species *Serritaenia testaceovaginata* (Fucíková, J.D. Hall, J.R. Johansen & R.L.Lowe) A.Busch & S.Hess 2021: 115.

Basionym: *Mesotaenium testaceovaginatum* Fucikova, J.D. Hall, J.R. Johansen & R.Lowe 2008: 31, pl. VI [6]: figs 6, 26-28.

LT: Fučíková et al., 2008: pl. VI, fig. 6;

Epitype: Permanent slide accession B 40 0001077 in Herbarium Berolinense (Berlin, Germany).

### Order Zygnematales

Bessey emend. S. Hess & J. de Vries, Bessey 1907, 9.

#### Emended diagnosis

Comprises unicells and unbranched and uniseriate filaments with smooth side walls, cells with asteroidal, plate-or ribbon-like chloroplasts and simple cell walls (no pores and ornamentations), phylogenetically closely related to strains SAG 698-1a (Genbank transcriptome shotgun assembly GFYA00000000).

#### Type family

Zygnemataceae Kützing 1843: 179, 274.

#### Type genus

Zygnema C. Agardh, 1817, nom. et typ. cons.

#### Type species

*Zygnema cruciatum* (Vaucher) C.Agardh 1817: 102.

#### Comment

Currently this order includes the genera *Cylindrocystis, Mesotaenium* (current assumption, pending discovery of type species), *Mougeotia, Zygnema*, and *Zygnemopsis*. No culture is available from the type species of *Zygnema*.

### Order Desmidiales

Bessey emend. Hess & de Vries, Bessey 1910, 88.

#### Emended diagnosis

Comprises unicells and chain-like filaments. Cell walls and morphologies of diverse complexity, including the “placoderm desmids” with cell wall pores, ornamentations and clear isthmus, and species with smooth cell walls and without isthmus. Phylogenetically closely related to strain *Desmidium aptogonum* (RNA-seq ERX2100155).

#### Type family

Desmidiaceae Ralfs 1848: 49.

#### Type genus

*Desmidium* C.Agardh ex Ralfs, 1848.

LT: *Desmidium swartzii* C.Agardh ex Ralfs 1848: 61, pl. IV [4]: figs a-f.

#### Comment

Currently this order includes the genera *Bambusina, Closterium, Cosmarium, Cosmocladium, Desmidium, Euastrum, Micrasterias, Netrium, Nucleotaenium, Onychonema, Penium, Phymatodocis, Planotaenium, Pleurotaenium, Staurastrum, Staurodesmus, Xanthidium*, and some more. No culture is available from the type species of *Desmidium*.

### Order Spirogyrales

Hess & de Vries ordo nov., *non* Clements 1909

#### Emended diagnosis

Comprises filaments with smooth side walls, cells with one or more helical chloroplast and smooth cell walls without pores or ornamentation. Phylogenetically closely related to strain *Spirogyra pratensis* strain MZCH10213 (RNA-seq data: NCBI BioProject PRJNA543475, TSA GICF00000000).

#### Type family

Spirogyraceae Blackman & Tansley 1902: 214.

#### Type genus

*Spirogyra* Link, 1820, nom. cons.

LT: *Spirogyra porticalis* (O.F.Müller) Dumortier

#### Comment

Currently this order only includes the genus *Spirogyra*. Probably the closely related genus *Sirogonium* Kützing also belongs to this order, but this needs to be confirmed by phylogenomic studies. No culture is available from the type species of *Spirogyra*. The order Spirogyrales Clements (1909) comprises exclusively zygomycetes, and not green algae, this order is not accepted by the mycologist community. Therefore, we reuse the name Spirogyrales because no priority rule above family level occurs according to the ICN.

## ACKNOWLEDGEMENTS

This work was funded by the German Research Foundation grants 283693520 (Research Fellowship) and 417585753 (Emmy Noether Programme) both to S.H., grant 440231723 (VR 132/4-1) within the framework of the Priority Programme “MAdLand – Molecular Adaptation to Land: Plant Evolution to Change” (SPP 2237) to J.d.V, and grant 410739858 in the frame of the project CharMod to K.v.S. J.d.V. further thanks the European Research Council for funding under the European Union’s Horizon 2020 research and innovation programme (Grant Agreement No. 852725; ERC-StG “TerreStriAL”). We thank Richard McCourt (Drexel University) and the Herbarium of the Academy of Natural Sciences of Philadelphia (PH) for destructive sampling of material from *Mesogerron fluitans*, and Elke Woelken (Universität Hamburg) for excellent support in electron microscopy.

## AUTHOR CONTRIBUTIONS

Conceptualization: S.H., J.d.V.;

Investigation: S.H., S.W., A.B., I.I., S.d.V., H.B., K.v.S., J.d.V.;

Writing - Original Draft: S.H., I.I., and J.d.V.;

Writing - Review & Editing: All authors;

Visualization: S.H., A.B., I.I., and J.d.V.;

Funding acquisition: S.H.; J.d.V.

## DECLARATION OF INTERESTS

The authors declare no competing interests.

## METHODS

### Resource table

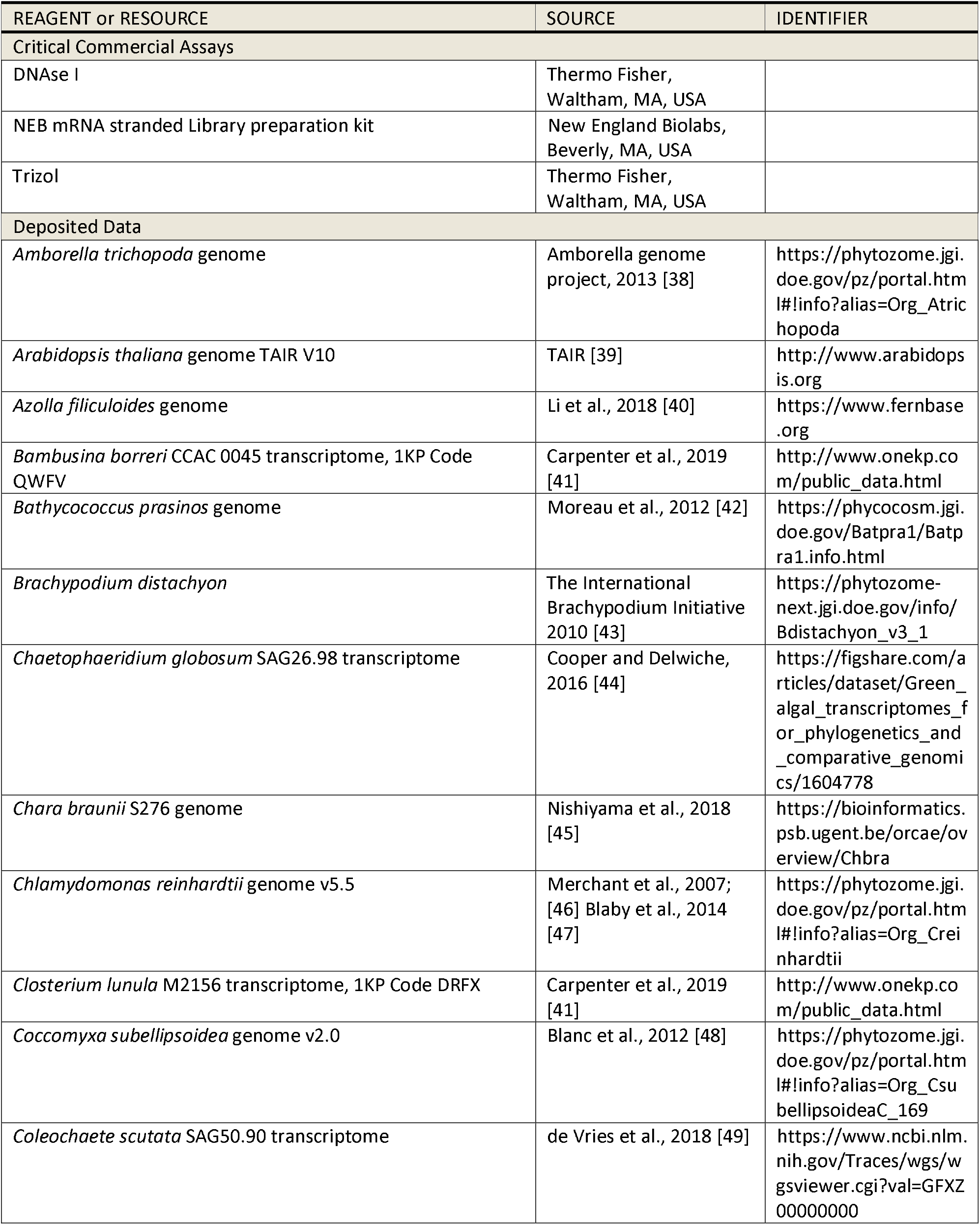
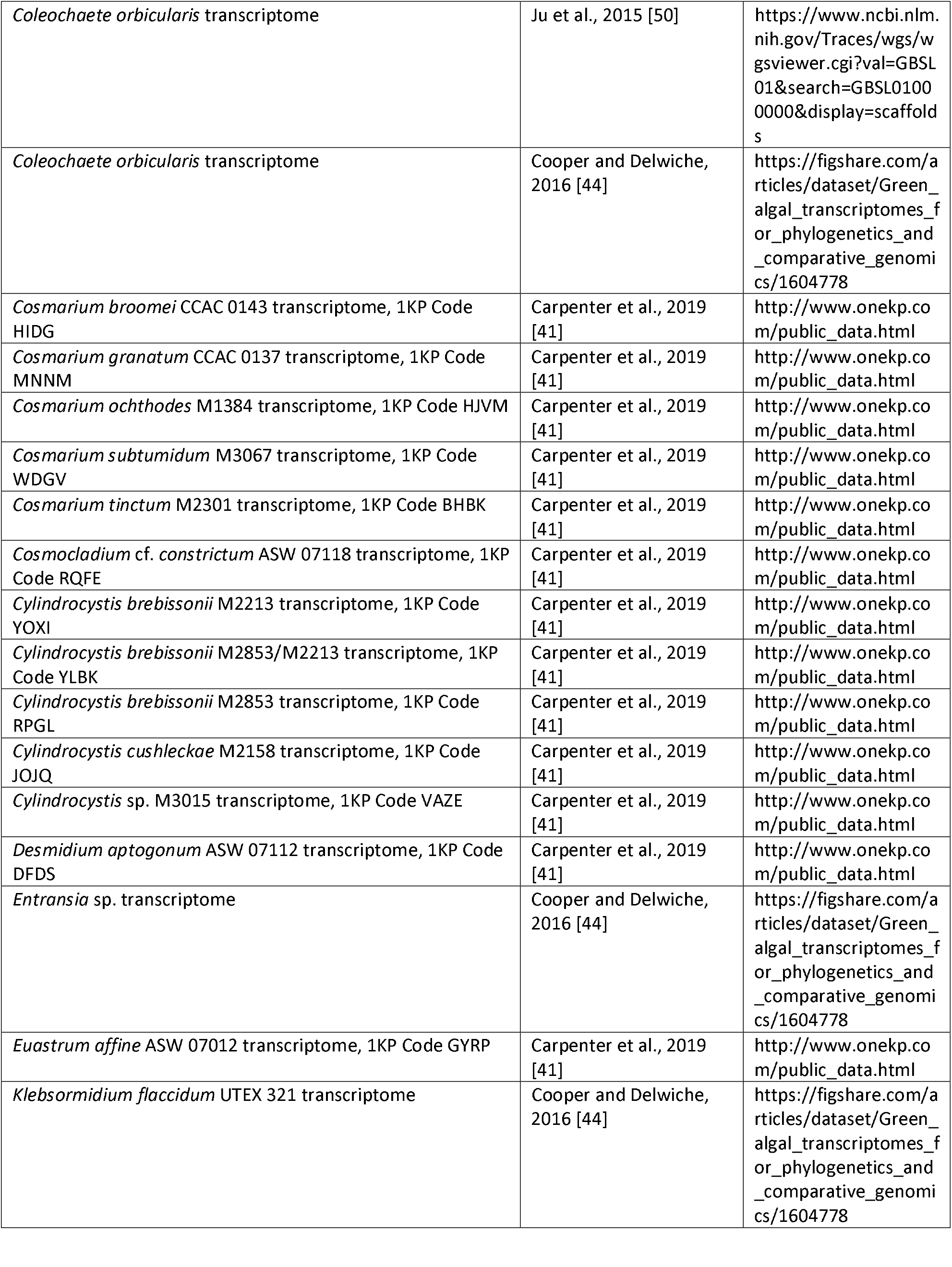
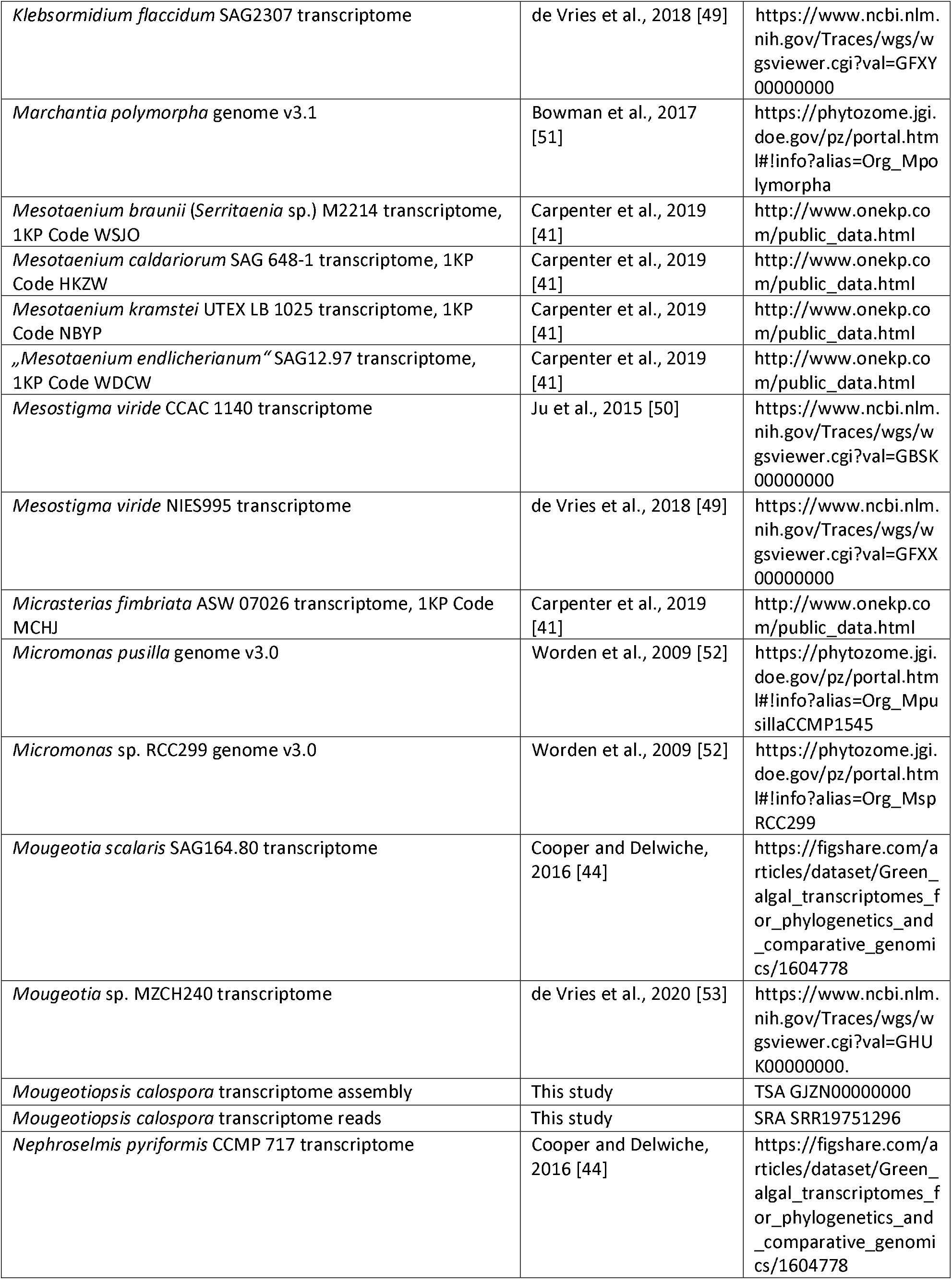
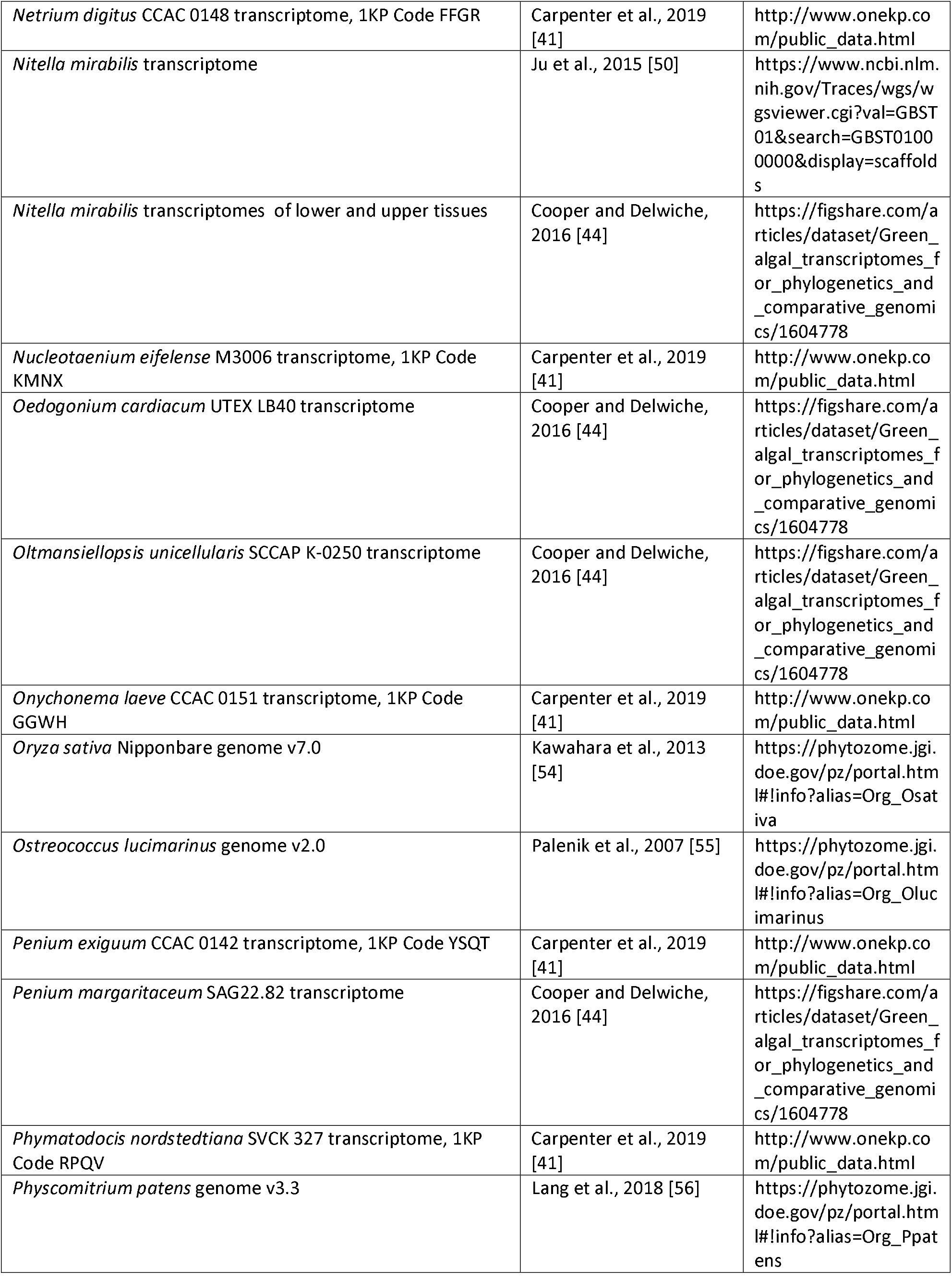
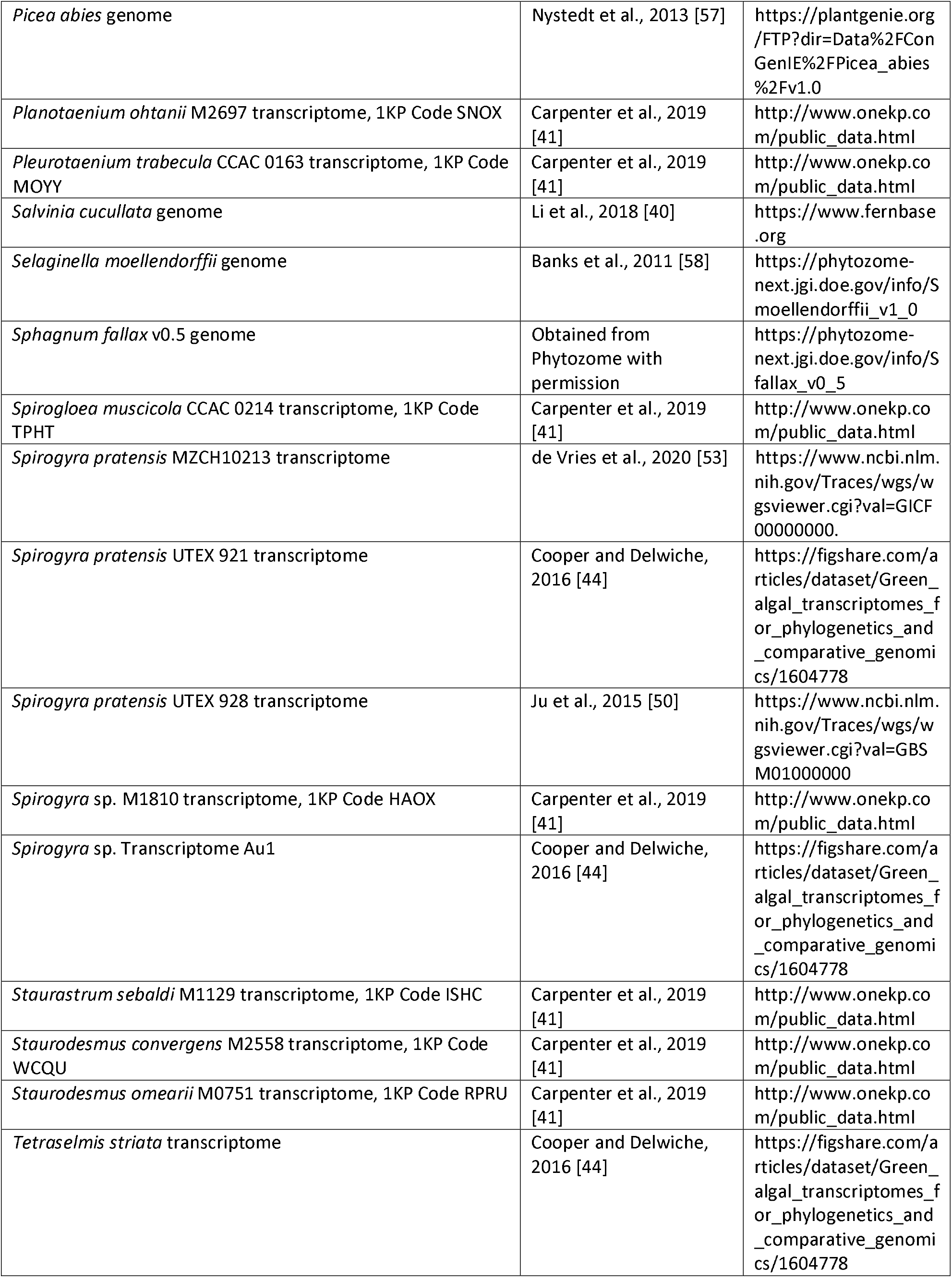
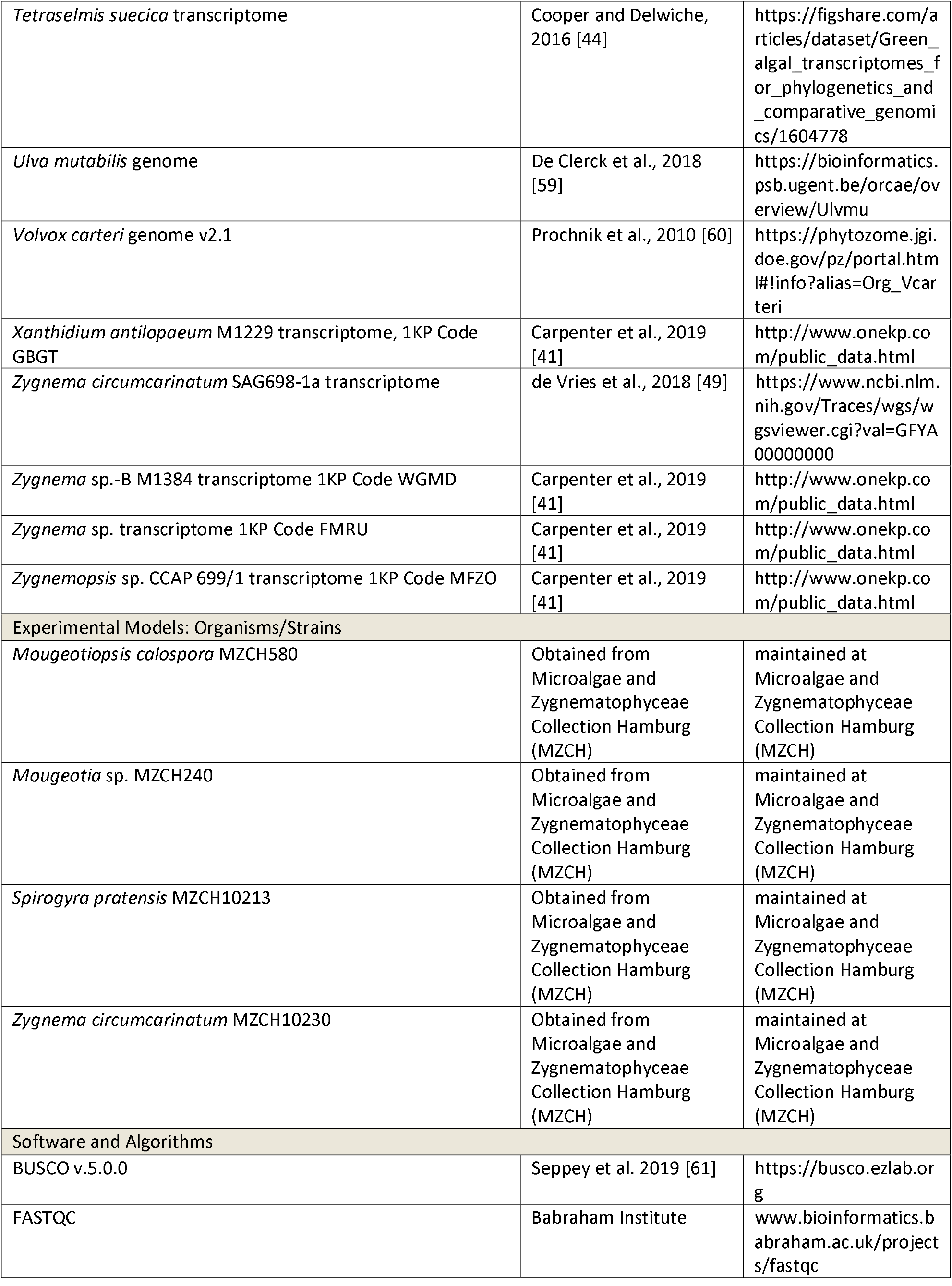
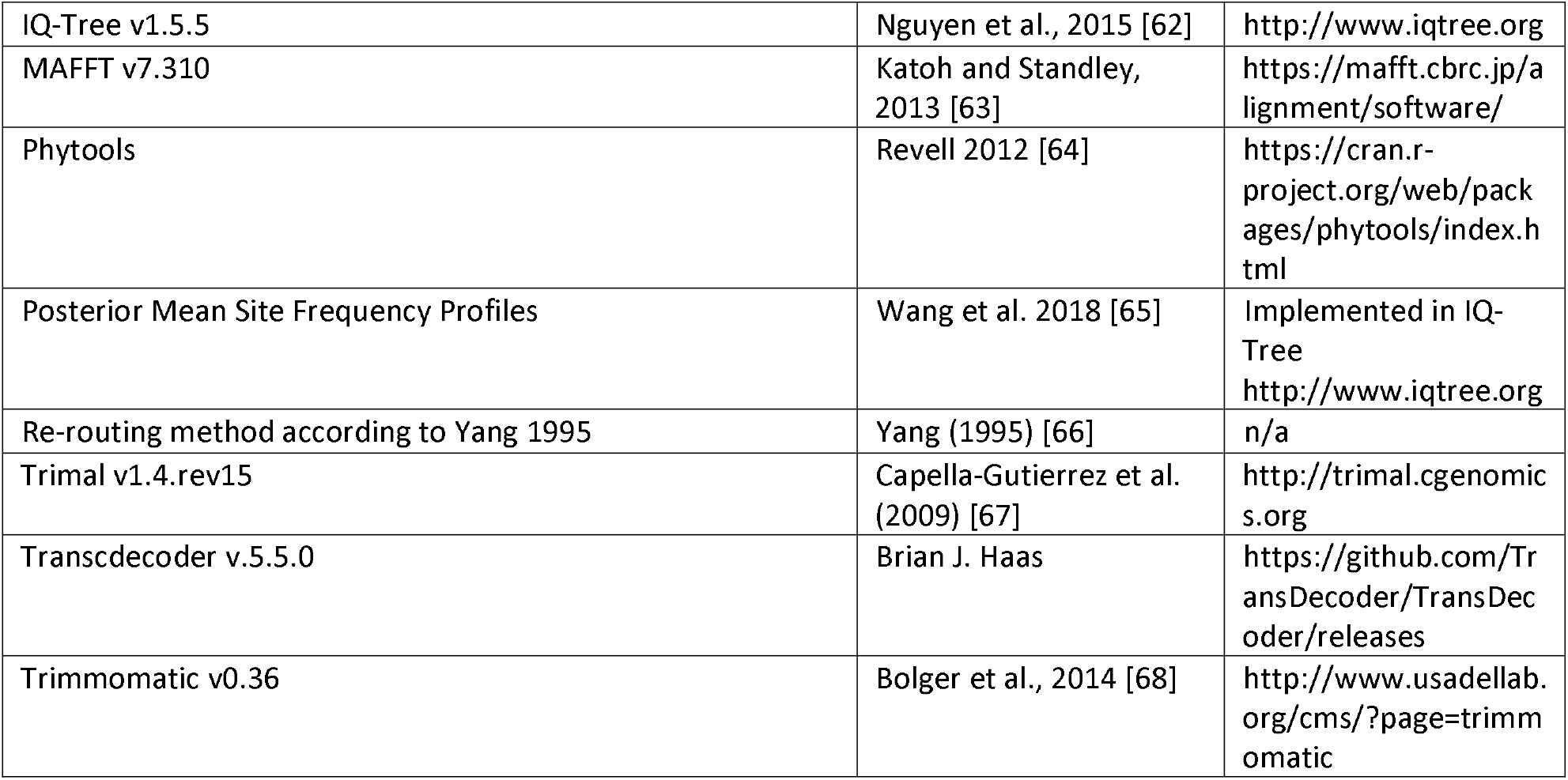

### Resource availability

#### Lead contact

Further information and requests for resources and reagents should be directed to and will be fulfilled by the lead contact, Jan de Vries.

### Materials availability

This study did not generate new unique reagents.

### Data and code availability

- RNA-seq data have been deposited at the NCBI under the BioProject accession PRJNA849386 and the Sequence Read Archive (SRA) under the accession SRR19751296; all data are publicly available as of the date of publication. Accession numbers are additionally listed in the key resources table.
- A transcriptome assembly has been deposited at NCBI Transcriptome Shotgun Assembly Sequence Database (TSA) under the accession GJZN00000000. The version described in this paper is the first version, GJZN01000000. The assembly is publicly available as of the date of publication. The accession number is additionally listed in the key resources table.
- No original code was used; all computational analyses were performed with published tools and are cited in the Methods section.

### Organisms

#### Algal strains

*Mougeotiopsis calospora* (strain MZCH580), *Mougeotia* sp. (MZCH240), *Spirogyra pratensis* (strain MZCH20213) and *Zygnema circumcarinatum* (MZCH10230) were obtained from the Microalgae and Zygnematophyceae Collection Hamburg (MZCH) [69, 70] and grown in WHM medium [71] or Waris-H medium [72] at 20°C and under fluorescent light or white LEDs (30-50 μmol photons m^-2^ s^-1^; 16h:8h light-dark cycle), if not stated otherwise in the experimental details (see below).

## Method details

### Light Microscopy, time-lapse photography, and confocal imaging

High-resolution imaging of *Mougeotiopsis calospora* was done with the Zeiss IM35 inverted microscope (Carl Zeiss, Oberkochen, Germany) equipped with the objective lens Planapochromat 63×/1.4, electronic flash, and the Canon EOS 6D digital single-lens reflex camera (Canon, Tokyo, Japan). Time lapse imaging was performed on a Leica DM5000B microscope (Leica Microsystems Wetzlar GmbH, Wetzlar, Germany) controlled by the Micromanager software at six frames per minute, shown as 10 FPS. Colour balance and contrast of micrographs were adjusted with Photoshop CS4 (Adobe Inc., CA, USA). Confocal laser scanning microscopy was done with a Leica TCS SPE system (SP5) and the Leica LCS software (Leica Microsystems Wetzlar GmbH, Wetzlar, Germany). Chlorophyll was excited with a wavelength of 635 nm and the emission of 646–782 nm was recorded. Confocal z-stacks were processed and converted to three-dimensional data with the image processing package Fiji [73].

### Transmission electron microscopy

Algal filaments were fixed with 2 % glutaraldehyde in 75 mM cacodylate buffer (pH 7.0) for 1 h at RT, rinsed with 75 mM cacodylate buffer, and postfixed with 1 % osmium tetroxide in 75 mM cacodylate buffer overnight at 4 °C. After rinsing in cacodylate buffer, the samples were dehydrated in a graded acetone series and embedded according to Spurr [74]. The resulting TEM blocks were sectioned on an Ultracut E ultramicrotome (Leica-Reichert-Jung, Vienna, AU), stained with 2 % uranyl acetate and 2 % lead citrate. Sections were then examined with the LEO 906E transmission electron microscope (LEO, Oberkochen, Germany) and imaged with a MultiScan Typ 794 CCD camera and the Digital Micrograph 3.4.4 software (both Gatan Inc., Pleasanton, USA).

### RNA isolation, sequencing and phylogenomics

For the isolation of total RNA, *Mougeotiopsis calospora* was grown on a modified freshwater F/2 medium [75] with 1% agar at 22°C. An LED light source provided photosynthetically active radiance at 120 μmol photons*m^-2^*s^-1^ under a 12:12 h light/dark photocycle. Harvesting, RNA extraction and transcriptome sequencing was carried out as described by de Vries et al. [53]. In brief, filaments of a growing algal culture were harvested and directly transferred into Trizol (Thermo Fisher, Waltham, MA, USA). The algal sample was homogenized using a Tenbroek tissue homogenizer and all following steps were performed in accordance to the manufacturer’s instructions. To remove possible residual DNA, RNA samples were treated with DNAse I (Thermo Fisher). Adequate RNA quality was verified using a formamide agarose gel. Samples were shipped on dry ice to Genome Québec (Montreal, Canada), where additional RNA quantification and quality assessments were performed using a Bioanalyzer (Agilent Technologies Inc., Santa Clara, CA, USA). Library construction was performed using the NEB mRNA stranded Library preparation kit (New England Biolabs, Beverly, MA, USA). Sequencing of the libraries was carried out on the NovaSeq 6000 (Illumina), yielding 28188133 paired end reads of 101 base pairs in length. Quality of the reads was assessed using FastQC version 0.11.7. Reads were trimmed using Trimmomatic version 0.36 [68], applying settings for quality trimming and adapter removal (ILLUMINACLIP:Adapters.fa:2:30:10:2:TRUE HEADCROP:10 TRAILING:3 SLIDINGWINDOW:4:20 MINLEN:36). The transcriptome was assembled de novo with Trinity. Transcriptome completeness was assessed with BUSCO v.5.0.0 using the viridiplantae_odb10 database in the transcriptome mode. Open reading frames (ORFs) were predicted with Transdecoder v.5.5.0.

We downloaded 83 transcriptomes and genomes of Streptophyta and Chlorophyta (see Resource table). Using a previously constructed phylogenomic dataset, we searched the selected sequencing data for orthologs of the 351 highly conserved proteins [76]. After alignment and trimming using MAFFT v7.310 [63] and trimal v1.4.rev15 [67], careful inspection of single-protein phylogenies estimated with IQ-TREE v1.5.5 under the LG4X model was undertaken to remove contaminants and paralogs. Once the data set was refined, orthologs with too much missing data were removed and we estimated a maximum likelihood phylogeny based on the concatenated alignment of a final set of 326 translated proteins (cumulative maximum of 115,424 sites); the final set of proteins/protein-coding genes was: AAP, ABHD13, Actin, ADK2, AGB1, AGX, AKTIP, ALG11, ALIS1, AMP2B, AOAH, AP1S2, AP3M1, AP3S1, AP4M, AP4S1, APBLC, ar21, arf3, ARL6, ARP2, ARP3, arpc1, ARPC4, ATEHD2, ATG2, atp6, ATP6V0A1, ATP6V0D1, ATPDIL14, ATSAR2, Atub, BAT1, Btub, C16orf80, C22orf28, C3H4, calr, capz, CC1, CCDC113, CCDC37, CCDC40, CCDC65, cct-A, cct-B, cct-D, cct-E, cct-G, cct-N, cct-T, cct-Z, CDK5, CLAT, COP-beta, COPE, COPG2, COPS2, COPS6, COQ4-mito, CORO1C, crfg, CRNL1, CS, CTP, D2HGDH-mito, DCAF13, DHSA1, DHSB3, DHYS, DIMT1L, DNAI2, DNAJ, DNAL1, DNM, DPP3, DRG2, ECHM, EF2, EFG-mito, EFTUD1, EIF3B, EIF3C, EIF3I, EIF4A3, EIF4E, ERLIN1, ETFA, FA2H, FAH, FAM18B, FAM96B, FAM, fh, fibri, FOLD, fpps, FTSJ1, GAS8, GCST, gdi2, GDI, glcn, GLGB2, GMPP3, gnb2l, gnbpa, GNL2, grc5, GRWD1, GSS, Gtub, H2A, H2B, h3, h4, HDDC2, HGO, HM13, hmt1, HSP70C, hsp70mt, HSP90, HYOU1, if2b, if2g, if2p, if6, IFT46, IFT57, IFT88, IMB1, IMP4, ino1, IP5PD, IPO4, IPO5, KARS, KDELR2, l10a, l12e-D, LRRC48, mat, mcm-A, mcm-B, mcm-C, mcm-D, mcm-E, metap2, METTL1, MLST8, MMAA-mito, mra1, MTHFR, MTLPD2, MYG1, NAA15, NAE1, NAPA, ndf1, NDUFV2-mito, NFS1-mito, NMD3, NMT1, NOP5A, NSA2, nsf1-C, nsf1-E, nsf1-G, nsf1-H, nsf1-I, nsf1-J, nsf1-K, nsf1-L, nsf1-M, nsf2-A, nsf2-F, ODB2, ODBA, ODBB, ODO2A, ODPA2, ODPB, oplah, orf2, osgep, PABPC4, pace2-A, pace2B, Pace2C, pace5, PCY2, PELO, PGM2, PIK3C3, PLS3, PMM2, PMPCB, PPP2R3, PPP2R5C, PPX2, PR19A, PSD11, PSD7, psma-A, psma-B, psma-C, psma-E, psma-F, psma-G, psma-H, psma-J, psmb-K, psmb-L, psmb-M, psmb-N, PSMD12, PSMD6, psmd, PURA, PYGB, rac, rad23, Rad51A, ran, RBX1, rf1, rla2a, rla2b, RPAC1, RPF1, rpl11, rpl12, Rpl13A, Rpl13e, Rpl14e, Rpl15, rpl17, Rpl18, rpl19, rpl20, rpl21, Rpl24A, rpl26, rpl27, Rpl2, rpl30, rpl31, rpl32, rpl33, rpl35, Rpl3, rpl43, rpl44, Rpl4b, Rpl5, rpl6, Rpl7a, rpl9, RPN1B, rpo-A, rpo-B, rpo-C, RPPK, rppO, rps10, rps11, rps12, rps14, rps15, rps16, rps17, rps18, rps20, rps23, rps26, rps27, rps2, rps3, rps4, rps5, rps6, rps8, RPTOR, RRAGD, RRM1, s15a, s15p, sap40, SCO1-mito, SCSB, SEC23, SF3B2, SND1, SPTLC1, sra, srp54, STXBP1, suca, SYGM1, SYNJ, tfiid, TM9SF1, TMS, topo1, trs, UBA3, ubc, UBE12, UBE2J2, Ubq, VAPA, VARS, vata, vatb, vatc, vate, VBP1, VPS18, VPS26B, WBSCR22, WD66, wd, wrs, xpb, YKT6. This tree was used as a guide to infer the final phylogeny under the LG+PMSF(C60)+F+Γ model [64] of evolution. Bootstrap analysis was conducted with 100 nonparametric bootstrap replicates using this model.

### Ancestral character state reconstruction

Ancestral character state reconstruction was performed with Phytools (Revell 2012 [64]), which implements Yang’s [66] re-rooting method to infer marginal ancestral state estimates for the internal nodes in the tree (Figure 3). We performed two independent analyses assuming 2-, and 4-character states in order to understand the effect of character coding on the inferred ancestral character states. The 2-state model used (1) unicellular and (2) multicellular *sensu lato* (filamentous or multicellular); the 4-state model differentiated between (2) *bona fide* filamentous algae excluding desmids, (3) chain-like filamentous desmids, and (4) multicellular *sensu stricto* (embryophytes, Coleochaetophyceae, Charophyceae, *Volvox, Ulva*). All models assumed unordered states (equal rates of change).

## SUPPLEMENTAL VIDEO

**Video S1: Cytokinesis in Mougeotiopsis calospora: centripetal cell division**. The movie shows that cytokinesis is dominated by centripetal cleavage. Note the rotating chloroplasts and the migrating nuclei. Imaging was performed at six frames per minute, shown as 10 FPS. Related to Figure 1.

**Video S2: Cytokinesis in Mougeotiopsis calospora**. A second movie to illustrate the different phenotypes of cell division; see description for Video S1. Related to Figure 1.

---

**Figure S1:**
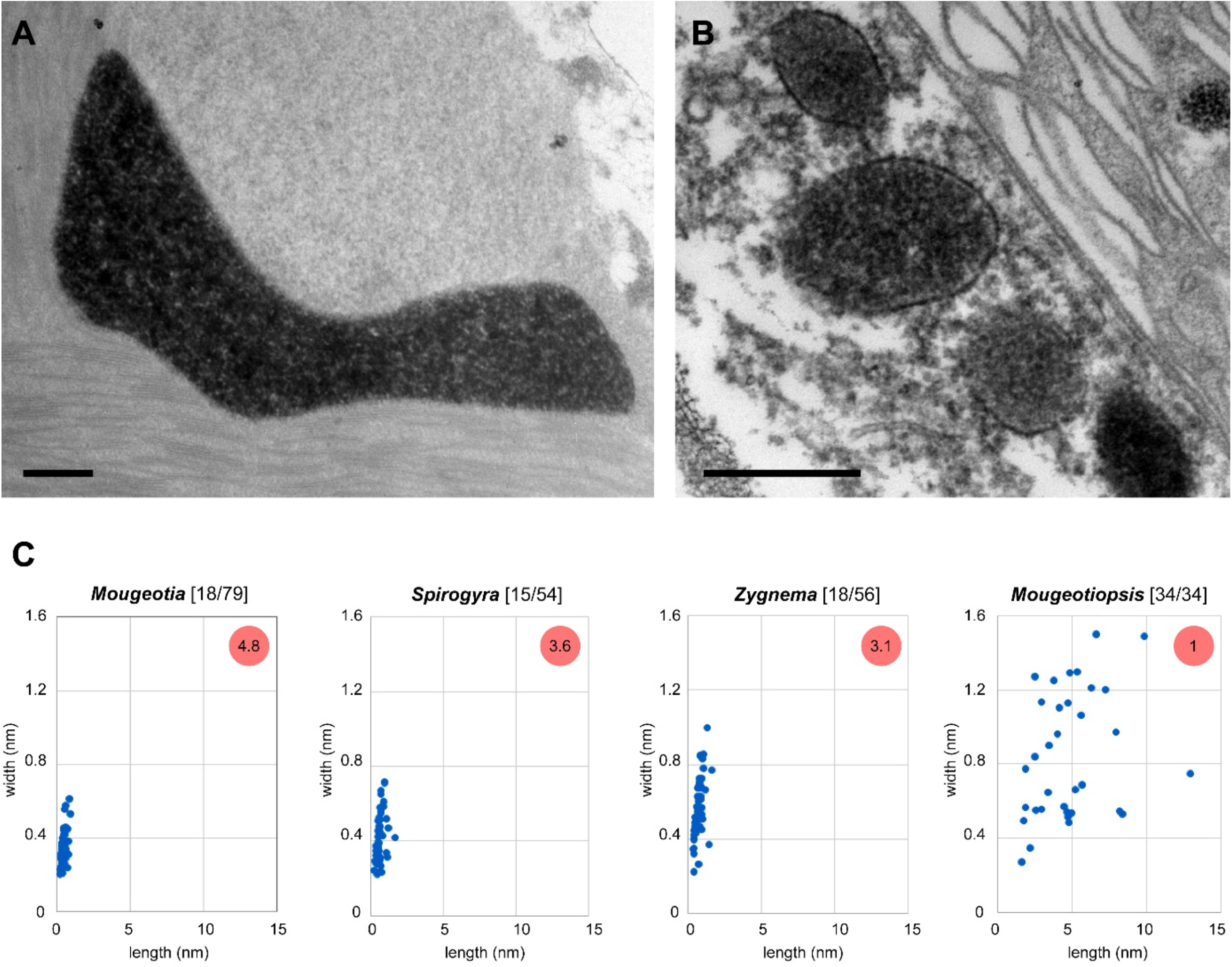
Peroxisomes of filamentous zygnematophytes, related to Figure 1. **A:** Transmission electron micrograph of a DAB-stained peroxisome of *Mougeotiopsis calospora*, strain MZCH580. **B:** Transmission electron micrograph of DAB-stained peroxisomes of *Mougeotia* sp., strain MZCH240. **C:** Sizes of peroxisome sections of four filamentous zygnematophytes as measured in transmission electron micrographs. The number of analysed cells/peroxisomes are shown in square brackets, and the average number of peroxisome cross sections per cell in the red circles. Scale bars, 0.5 μm.

**Figure S2:**
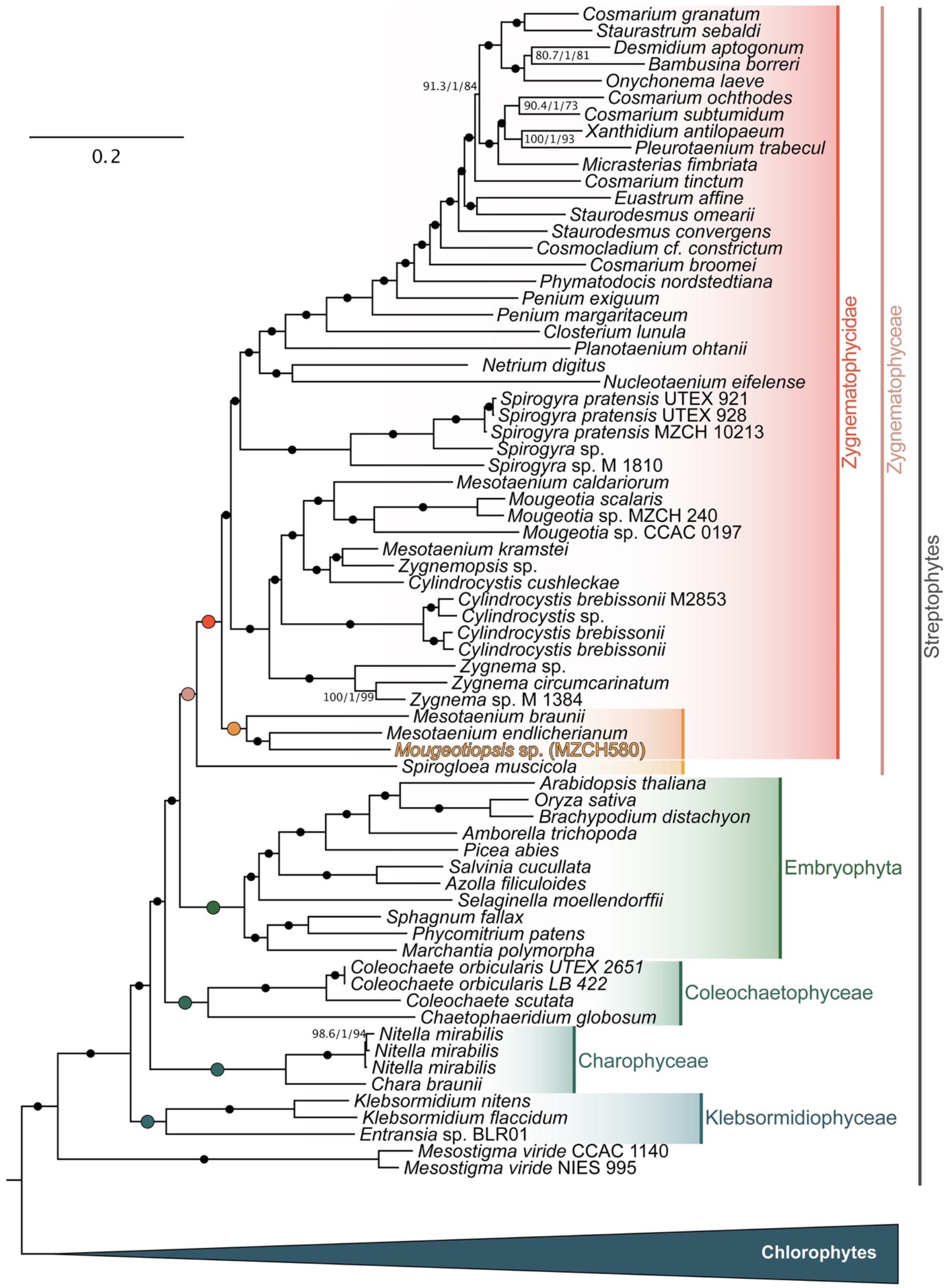
Multigene phylogeny of 84 Viridiplantae, related to Figure 2. Phylogenomic tree that shows the relationship of all streptophyte species analysed; the tree was rooted with the clade of chlorophytes. Scale bar, 0.2 substitutions per site.

**Figure S3:**
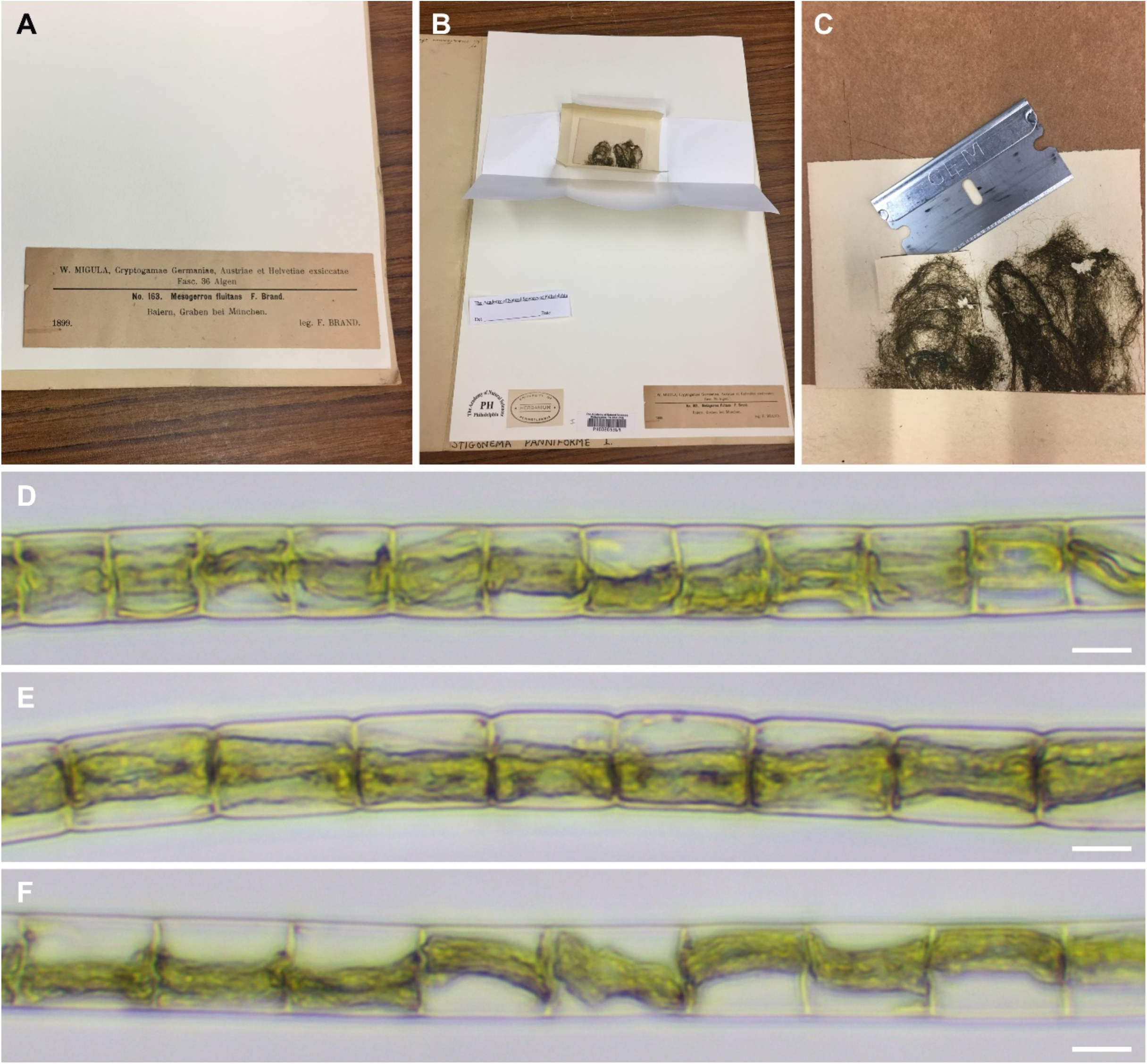
Destructive sampling of *Mesogerron fluitans* collected by Brand and morphological characteristics of the material, related to Figure 1. **A and B:** Specimen in the Herbarium of the Academy of Natural Sciences of Philadelphia (PH). **C:** Removal of dried algal material. **D–F:** Rehydrated algal filaments of the sample. Note the varying cell length and the chloroplast morphology resembling that of strain MZCH580. Images in A–C: courtesy of Richard McCourt. Scale bars, 10 μm.

## Notes

### Competing Interest Statement

The authors have declared no competing interest.

